# Deciphering the mRNA-lncRNA-miRNA interaction landscape in human pluripotency

**DOI:** 10.1101/2022.04.12.488044

**Authors:** Arindam Ghosh, Anup Som

**Author notes:** Corresponding Author: Anup Som.

## Abstract

Human embryonic stem cells offer a huge potential for the study of early human development and for application in biomedical sciences. Growing evidence has shown that besides the protein coding genes, the non-coding elements of the human genome play a crucial role in maintaining its property of self renewal and in cell fate determination. However, a clear understanding of this regulatory mechanism and the landscape of interactions between the coding and non-coding elements was still lacking. To fill in this void, we use transcriptomic data from RNA-seq and small RNA-seq experiments to reconstruct the core pluripotency circuitry involving mRNAs, lncRNAs and miRNAs. The overall interaction landscape revealed an alternate circuit for the maintenance of pluripotency devoid of the classic pluripotency transcription factors NANOG, SOX2 and POU5F1. We also identified networks specific to the naive and primed states of human pluripotency revealing a new set of transcriptomic markers that could not only be used to differentiate pluripotent state from non-pluripotent state but also to identify the intra-pluripotency state. The lncRNA DANT1 was found to be crucial in determination to the two pluripotency states as it formed a bridge between the naive and primed state specific pluripotency networks. Further, we also identified and computationally validated putative ceRNA mechanism involving DANT1, the miRNAs hsa-miR-30c-2-3p, hsa-miR-210-3p and hsa-let-7b-5p, and several key pluripotency related genes including PTPRZ1, SALL2, TOX3, ZNF695, and ZYG11A which warrants further experimental validation.

**HIGHLIGHTS:**

- Reconstructed the key MLMi circuitry underlying human pluripotency by combining RNA-Seq data and known interaction information.
- Identified an alternate pluripotency circuit devoid of classic pluripotency transcription factors NANOG, SOX2 and POU5F1.
- Predicted a new set of markers that can not only distinguish between pluripotent and non-pluripotent states but also identify the intra-pluripotency state.
- Identified novel putative function of lncRNA DANT1 as ceRNA in formation of naive and primed state pluripotency.
- DANT1 harbours binding sites for miRNAs hsa-miR-30c-2-3p, hsa-miR-210-3p and hsa-let-7b-5p.

## 1. INTRODUCTION

Pluripotency is the ability of a cell to differentiate into cells of any of the three germ layers (i.e., endoderm, mesoderm, and ectoderm). Besides, it also has the property of self-renewal that provides a unique opportunity to human pluripotent stem cells such as embryonic stem cells (ESCs) to be used for studying early human development and in therapeutic applications (De Los Angeles et al., 2015; Yamanaka, 2020). However, their applications have been limited due to the lack of complete understanding of the mechanisms of pluripotency. Initial efforts in this direction mainly focussed on the coding elements of the genome. As such, several key transcription factors, namely NANOG, POU5F1, SOX2, STAT3 and ZIC3, have been discovered that are critical to pluripotency (Boyer et al., 2005). They have also been used to induce pluripotency in somatic cells resulting in the introduction of induced pluripotent stem cells (Takahashi and Yamanaka, 2006). While the protein coding genes play a critical role in maintaining this property, it has also been shown to be dependent on several non-protein coding elements of the human genome such as lncRNAs and miRNAs (Leonardo et al., 2012; Lu et al., 2021; Rosa and Brivanlou, 2013). They primarily aid in modulating the expression of the protein coding genes. Across biological systems, the importance of these non-protein coding elements, which was previously considered as junk, is being increasingly realised. In context to pluripotency, the role of the ncRNAs have mostly been explored independently (Mirzadeh Azad et al., 2021) and there are limited studies that try to provide a detailed interaction landscape between the coding and non-coding elements. In this study, we reconstruct the core circuit consisting of mRNAs, lncRNAs and miRNAs (MLMi network) that might be involved in maintaining pluripotency in humans. Transcriptomic data from RNA-seq and small RNA-seq, measuring the expression of mRNAs and lncRNAs, and miRNAs respectively were used for the purpose. We used a two stage approach for finding the core links between all the three RNA types. In the first stage, we created a base network of the mRNAs and the lncRNA using the weighted gene co-expression network analysis (WGCNA) approach and extracted the core network. In the second stage, we integrated the miRNAs into this core mRNA-lncRNA network by retrieving interaction from the miRNet database. Analysis of this network revealed a possible alternate circuitry for maintaining pluripotency that does not include the classic pluripotency transcription factors NANOG, POU5F1 and SOX2. Further, utilising previously identified naive and primed state specific protein coding genes, we derived the MLMi network specific to the two states. Analysis of these networks re-highlighted the role of some well-known pluripotency related genes besides suggesting some new markers including the lncRNA DANT1 that might be acting as a competing endogenous RNA (ceRNA) for regulating the expression of several mRNAs. Our analysis also revealed putative novel interactions between DANT1 and the miRNAs hsa-miR-30c-2-3p, hsa-miR-210-3p and hsa-let-7b-5p that complete the ceRNA circuit.

## 2. MATERIALS AND METHODS

### 2.1. Data retrieval

The RNA sequencing data for pluripotent and non-pluripotent samples were retrieved from the European Nucleotide Archive (ENA). A total of 25 pluripotent and 20 non-pluripotent samples from two datasets were included for mRNA and lncRNA expression profiling while 10 pluripotent and 9 non-pluripotent samples from another three datasets were included for miRNA expression profiling (Table 1). In each case, the datasets were chosen such that they contained both pluripotent and non-pluripotent samples. Further, only the datasets in which samples were enriched by removal of ribosomal RNAs included for profiling of mRNA and lncRNA expression (Supplementary File 1).

**Table 1:** Summary of data included in the study

| Accession number | Number of pluripotent samples | Number of non-pluripotent samples | Reference |
| --- | --- | --- | --- |
| **RNA-Seq** |  |  |  |
| PRJNA484413 | 4 | 4 | (Spencer et al., 2020) |
| PRJNA270993 | 21 | 16 | (Szabo et al., 2015) |
| **miRNA-Seq** |  |  |  |
| PRJNA282001 | 4 | 4 | (Jönsson et al., 2015) |
| PRJNA162685 | 3 | 2 | (Hu et al., 2012) |
| PRJNA422248 | 3 | 3 | (Garate et al., 2018) |

### 2.2. Data processing

The quality of the raw reads was assessed using FastQC v0.11.5 toolkit (Andrews, 2010) followed by using Trimmomatic v0.36 (Bolger et al., 2014) to remove the adapters and low-quality reads. The trimmed reads of the mRNA-lncRNA datasets were then mapped to the reference genome GRCh38.p5 (Ensembl release 84) using HiSat2 v2.1.0 (Kim et al., 2015) while Bowtie2 (Langmead and Salzberg, 2012) was used for mapping the miRNA reads to the miRBase miRNA reference sequences (Kozomara et al., 2019). The reads that mapped to the reference genome were used to estimate the expression of genes using featureCounts v1.6.2 (Liao et al., 2014) and annotations from Ensembl release 84 (for mRNA and lncRNA) or miRBAse (for miRNA). The genes which did not have more than one count per million reads (1 CPM) in at least fifty percent of the samples were identified as low expressed genes and removed from further analysis. The raw counts obtained from featureCounts were normalised using DESeq2 (Love et al., 2014). The presence of batch effect was accounted for using the function removeBatchEffect() from the R package limma (Ritchie et al., 2015). DESeq2 was also used to identify the differentially expressed genes – the genes with log2FC > 1.0 and adjusted p-value (padj) < 0.05 were considered upregulated or overexpressed in pluripotent state while those with log2FC < −1.0 and padj < 0.05 were considered downregulated or underexpressed in pluripotent state.

### 2.3. MLMi network reconstruction

The reconstruction of the core MLMi network underlying human pluripotency was performed in two stages. In the first stage, the expression values for mRNA and lncRNA were used to create a weighted co-expression network using the R package “WGCNA” (Langfelder and Horvath, 2008). The adjacency matrix representing the network was created by first calculating the biweight midcorrelation from the variance stabilised gene counts between all possible gene pairs followed by raising it to a power β (soft-threshold) which was in turn chosen based on the criteria of approximating the scale-free topology of the network (Zhang and Horvath, 2005). The adjacency matrix was then used to generate the Topological Overlap Measure (TOM) matrix for average linkage hierarchical clustering to identify gene clusters or modules. In our analysis, we identified β = 14, corresponding to a scale-free topology fit index R2 = 0.7 using the pickSoftThreshold function in WGCNA (Figure 1a). The parameters deepSplit = 2 and minimumModuleSize = 30 were used for network construction and module detection. A unique colour was assigned to each module for easy reference in downstream analyses. To determine whether the identified modules were significantly associated with either of the two sample types, a binary matrix was generated describing the association of the samples with their respective traits (pluripotent = 0 and non-pluripotent = 1). This matrix was then used as an input to calculate the correlation and p-value with the sample types. The modules with correlation coefficient greater than 0.8 and p-value < 0.05 were selected for further analysis. The intra modular hubs were selected based on gene significance (GS) and module membership (MM) (|GS| > 0.8 & p.GS < 0.05 & |MM| > 0.8 & p.MM < 0.05). The sub-networks of identified intra-modular hubs were created by retaining interactions with TOM > 0.55. Following this, the miRNAs were integrated into the mRNA-lncRNA network by retrieving interactions from the miRNet database (Chang et al., 2020). The mRNAs and lncRNAs in the core mRNA-lncRNA network were queried to retrieve all miRNAs that targeted them and then filtered to retain only those that were differentially expressed between pluripotent and non-pluripotent samples. The final set of mRNA-miRNA and lncRNA-miRNA interactions were integrated into the mRNA-lncRNA core network using union mode in Cytoscape (Shannon et al., 2003). Cytoscape was then used for network visualisation and further analysis.

**Figure 1:**
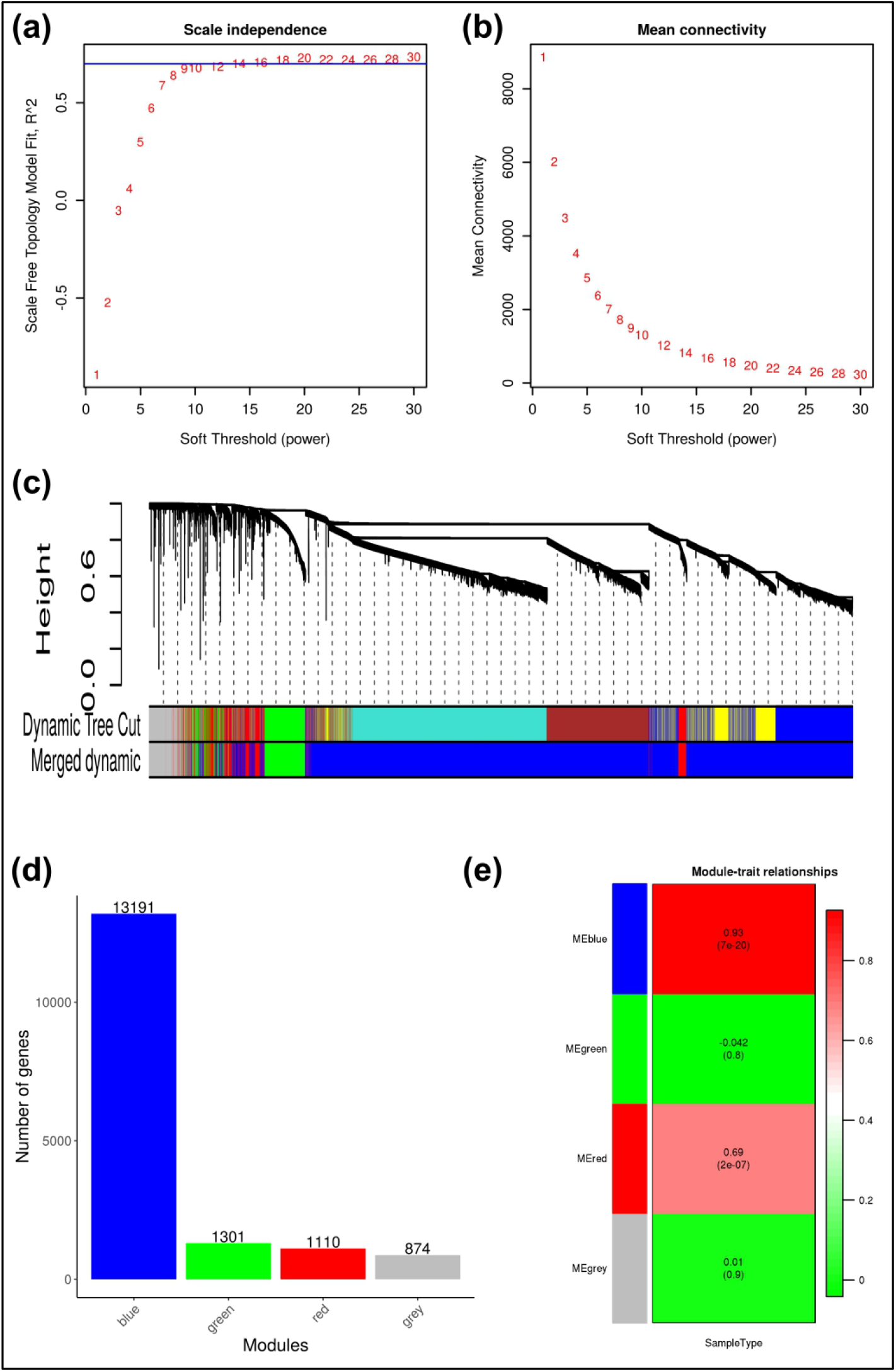
WGCNA for construction of mRNA-lncRNA network. (a,b) Determination of soft– thresholding power. (c) Hierarchical clustering tree of the entire set of mRNA and lncRNA. (d) Bar–plot depicting the number of genes in each module. (e) Module–trait relationship analysis. The values within each cell represent the correlation coefficient of the module–eigen with sample type (trait). The corresponding p–values for each module are mentioned within parentheses.

### 2.4. Enrichment analyses

Enrichment analyses of gene sets were carried out using the Enrichr web server (Chen et al., 2013). The input gene sets were compared against four gene set libraries in Enrichr – GO Biological Process 2021 for functional annotation, KEGG 2021 Human for pathway annotation, TRRUST Transcription Factors 2019 for identifying transcription factors that regulate the input gene set and CellMarker Augmented 2021 for identifying if the gene sets recapitulate any specific cell type. The enriched terms were sorted based on p-value and the top ten terms selected from each category.

### 2.5. Prediction of miRNA binding sites in lncRNA

The miRNA binding sites within lncRNAs were predicted from their sequences using the web-based tool rna22 (Miranda et al., 2006). The genomic sequences of the matured miRNA and the lncRNA were retrieved from RNA central and Ensembl respectively. We used the tool’s default parameters for identifying the heteroduplexes. The heteroduplexes with p-value < 0.01 were considered significant.

## 3. RESULTS AND DISCUSSIONS

### 3.1. Identification of differentially expressed mRNA, lncRNA and miRNA

The varying lengths of the RNAs are a major hurdle that impedes the correct measurement of gene expression in a single RNA-Seq experiment. It has been observed that the longer mRNAs and lncRNAs tend to be overrepresented leaving the shorter miRNAs under-represented. In order to avoid this, expression profiling of mRNA and lncRNA are done together while those of miRNA are done separately. Accordingly the differential gene expression analysis was performed separately. From differential gene expression analysis, we found that 5274 mRNAs (3098 upregulated, 2176 downregulated), 1025 lncRNA (357 upregulated, 668 downregulated), and 218 miRNA (108 upregulated, 110 downregulated) were differentially expressed between pluripotent and non-pluripotent samples (Supplementary File 2).

### 3.2. Reconstruction of the core MLMi network in pluripotency

We used a two stage approach in elucidating the core MLMi network involved in human pluripotency. In the first stage, we used the WGCNA approach to create a base network consisting of the mRNAs and lncRNAs. In the second stage, all the miRNAs that interacted with the mRNAs and lncRNAs in the mRNA-lncRNA network were retrieved from the miRNet database to create the integrated MLMi network. WGCNA was applied on a set of 16476 genes (mRNAs and lncRNAs) obtained after filtering the genes with low counts among the complete set and used to identify modules or groups of genes that worked in a coordinated manner for performing a specific biological function (Langfelder and Horvath, 2008). Three modules were identified with the blue module containing a majority of 13191 genes (Figure 1d). The other two modules – green and red, contained 1301 and 1110 genes respectively. The remaining of the 874 genes that could not be assigned to any of the clusters were added to the grey module. The modules that might be associated with the sample type i.e pluripotent or non-pluripotent, were identified by performing a module-trait analysis, which basically tries to find the correlation between the first principal component of the module’s gene expression (also called module eigen) with the sample traits. We observed that only the large blue module was significantly correlated (cor = 0.93, p-val = 7e-20) to the sample type (Figure 1e). Since the number of genes was too large to be taken forward for further analyses, we extracted the core or intra-modular hub network. This was done by filtering the nodes based on gene significance and module membership while the interactions were filtered based on topological overlap measure. The gene significance is a measure of biological significance of the gene with respect to a particular trait and is defined as the correlation between gene expression and sample trait. On the other hand, module membership defines how strongly the gene is associated with the module and is measured by correlating the gene expression with the module eigen. The topological overlap measure provides the strength of connection between two nodes with consideration to all other nodes in the network. We observed that the core mRNA-lncRNA network consisted of 432 nodes (375 mRNA and 57 lncRNA) connected by 5578 edges. Of the 432 nodes, 332 nodes were upregulated and 94 nodes were downregulated, while 6 nodes were not significantly differentially expressed but had log_2_FC close to 1. The miRNAs interacting with the 375 mRNA and 57 lncRNA were retrieved from the miRNet database and filtered to keep only those miRNA that were found to be differentially expressed between pluripotent and non-pluripotent samples. This resulted in 207 miRNAs connected to the mRNA and lncRNA with 4961 edges. Finally, the edges connecting mRNA and lncRNA (obtained from WGCNA), and mRNA and lncRNA with miRNA (obtained from miRNet) were merged using Cytoscape to obtain the final MLMi network, henceforth referred as PluriMLMiNet, containing 639 nodes (375 mRNA, 57 lncRNA and 207 miRNA) connected by 10539 edges. The topological parameters of the PluriMLMiNet are given in (Table 2) and the final network is provided as a separate cytoscape file (Supplementary File 3).

**Table 2:** Topological parameters of the networks

|  | **PluriMLMiNet** | **PluriMLMiHub**<br>**Net** | **NaiveMLMiNet** | **PrimedMLMi**<br>**Net** |
| --- | --- | --- | --- | --- |
| Number of nodes | 639 | 17 | 83 | 151 |
| Number of edges | 10539 | 56 | 144 | 431 |
| Avg. number of neighbours | 32.986 | 6.588 | 3.512 | 5.772 |
| Characteristic path length | 2.363 | 1.625 | 2.945 | 3.092 |
| Clustering coefficient | 0.049 | 0.149 | 0.075 | 0.043 |
| Connected components | 1 | 1 | 2 | 2 |
| Network density | 0.052 | 0.412 | 0.043 | 0.039 |
| Network diameter | 4 | 3 | 5 | 7 |
| Network radius | 3 | 2 | 3 | 4 |
| Network heterogeneity | 1.053 | 0.400 | 1.453 | 1.159 |
| Node degree exponent | 3.31 | 3.91 | 2.41 | 1.9 |

### 3.3. Enrichment analyses

Enrichment analysis is a key step towards understanding how a group of genes function. Since lncRNAs and miRNA execute their function by regulation of the mRNA, they are usually not well represented within ontology databases. Accordingly, we performed enrichment analysis of the mRNAs only. Enrichment analyses of the upregulated and downregulated mRNAs were carried out separately (Figure 2). We observed that the GO biological process terms tight junction assembly (GO:0120192), regulation of transcription, DNA-templated (GO:0006355) and endodermal cell fate specification (GO:0001714) to be significantly enriched among the upregulated mRNAs. The maintenance of hESC stemness is dependent on several intrinsic and extrinsic factors like transcriptional regulation and presence or absence of metabolites. Tight junction proteins play a pivotal role by regulating the flow of metabolites across the tight junctions formed surrounding stem cell clusters (Xing et al., 2020). Similarly, transcription regulation is a critical process that helps to maintain cellular identity. On the other hand, the terms related to positive regulation of programmed cell death (GO:0043068), extracellular matrix organisation (GO:0030198) and positive regulation of vesicle-mediated transport (GO:0060627) were among the enriched terms for downregulated mRNAs. Several previous studies have shown the link between programmed cell death and stem cells (Cheung et al., 2012; Ho et al., 2017; Hu et al., 2021). It might be particularly required during hESC differentiation to maintain the genomic integrity. While for structural formation and maintenance of the tissue, extracellular matrices play an important role. Enrichment analysis of the KEGG pathways revealed the upregulated mRNAs to be enriched for the signalling pathways regulating pluripotency of stem cells, while the downregulated mRNA were enriched for pathways in cancer. We also found the pathways related to cysteine and methionine metabolism, and nicotinate and nicotinamide metabolism among the top ten pathways enriched for upregulated mRNAs. They are primarily responsible for cell survival and regulation of differentiation of hESCs (Meng et al., 2018; Shiraki et al., 2014). In line with GO biological processes enrichment, we also saw the pathways related to cell adhesion molecules and tight junctions to be enriched for the upregulated mRNAs. Further, in order to identify the possible transcription factors that regulate the mRNAs in PluriMLMiNet, we performed transcription factor enrichment analysis. The upregulated mRNAs were primarily under the control of POU5F1 while the downregulated mRNA were found to be regulated by RUNX1. Among the top ten transcription factors, we can also see SOX2 and SALL4 which along with POU5F1 and NANOG are well-known for transcriptional regulation of pluripotency (Yeo and Ng, 2013). The transcription factor RUNX1, which was found to regulate the downregulated mRNAs, has been reported to be essential for differentiation of pluripotent stem cells to hematopoietic stem and progenitor cells (Teichweyde et al., 2017). Interestingly, we observed that none of the top ten transcription factors found to regulate the downregulated mRNAs are actually in PluriMLMiNet and must be remotely controlling the activation or inhibition of these genes. Finally, we performed enrichment analysis against cell markers to identify if these sets of mRNAs are able to recapitulate any specific cell types. We observed that the upregulated mRNAs were enriched for cell markers for several types of stem cells like pluripotent very small embryonic-like cells, gonocytes, ovarian germ stem cells, etc. While the downregulated mRNAs were mostly enriched for cell markers for differentiated cells like cardiac progenitor cell, cardiomyocyte, plasmacytoid dendritic cell, etc.

**Figure 2:**
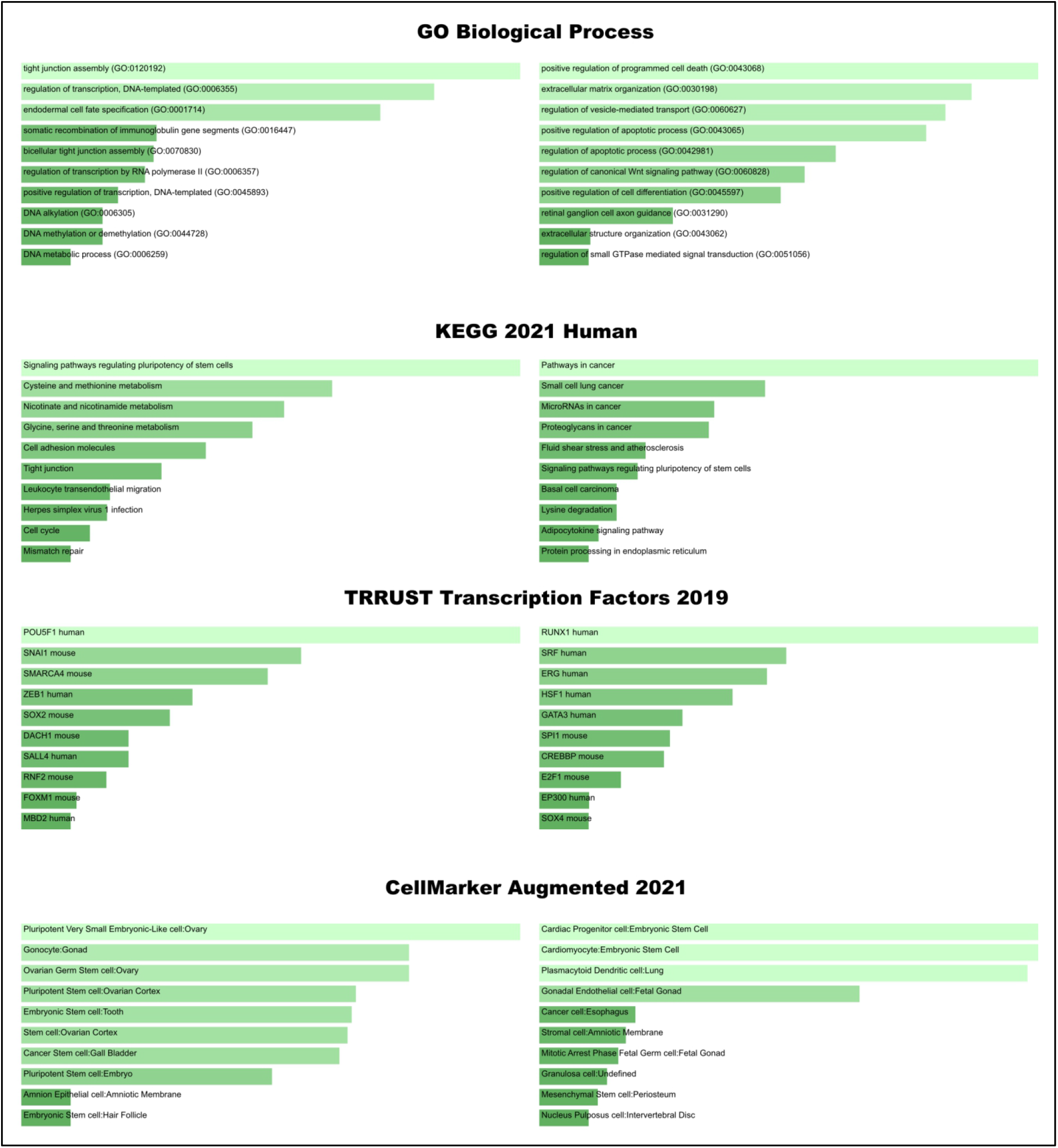
Enrichment analysis of GO biological process, KEGG human pathway, TRRUST transcription factors and cell markers. The left panel histograms are for upregulated mRNA while the right panel histograms are for downregulated mRNA. The colour of the histogram denotes the significance of the terms – lighter is more significant.

### 3.4. Identification of hub genes in PluriMLMiNet

Hub genes play a critical role in the dissemination of information within a network. Hub genes can be identified based on different centrality measures like degree, betweenness and closeness (Koschützki and Schreiber, 2008). The top twenty five genes in the network by degree centrality, betweenness centrality and closeness centrality were compared to identify the hub genes in the network. We observed 17 genes were common of which 14 were mRNA, two miRNA and one lncRNA (Figure 3). The 17 genes were used to extract the hub of the MLMi network or the PluriMLMiHubNet (Figure 4). Of the 14 mRNAs, 11 were upregulated and three were downregulated, while of the two miRNAs, hsa-miR-124-3p was upregulated and hsa-miR-155-5p was downregulated. The lone antisense lncRNA RP11-597D13.9 was downregulated. In one of our previous work, we had identified DPPA4, PRDM14, SCNN1A, and ZSCAN10 to be critical for pluripotency (Ghosh and Som, 2020). The genes DPPA4, PRDM14 and ZSCAN10 are known to be part of pathways related to transcriptional regulation of pluripotent stem cells. In the PluriMLMiHubNet, we observed that the lncRNA RP11-597D13.9, which has previously been associated with cancer progression (Zhan et al., 2018), interacts with three of these pluripotency critical genes – PRDM14, SCNN1A, and ZSCAN10, suggesting that it must also have an important regulatory mechanism in pluripotency. Other mRNAs in the hub network that have been linked to pluripotency include CDH1, GLDC, HOXB4, NFIA, PLD1, PMAIP1, and SEPHS1. Downregulation of CDH1 has been reported to result in loss of stem cell properties (Liu et al., 2009). GLDC is a metabolic regulator of pluripotent stem cells and is particularly involved in regulation of glycolysis (Kang et al., 2019).

**Figure 3:**
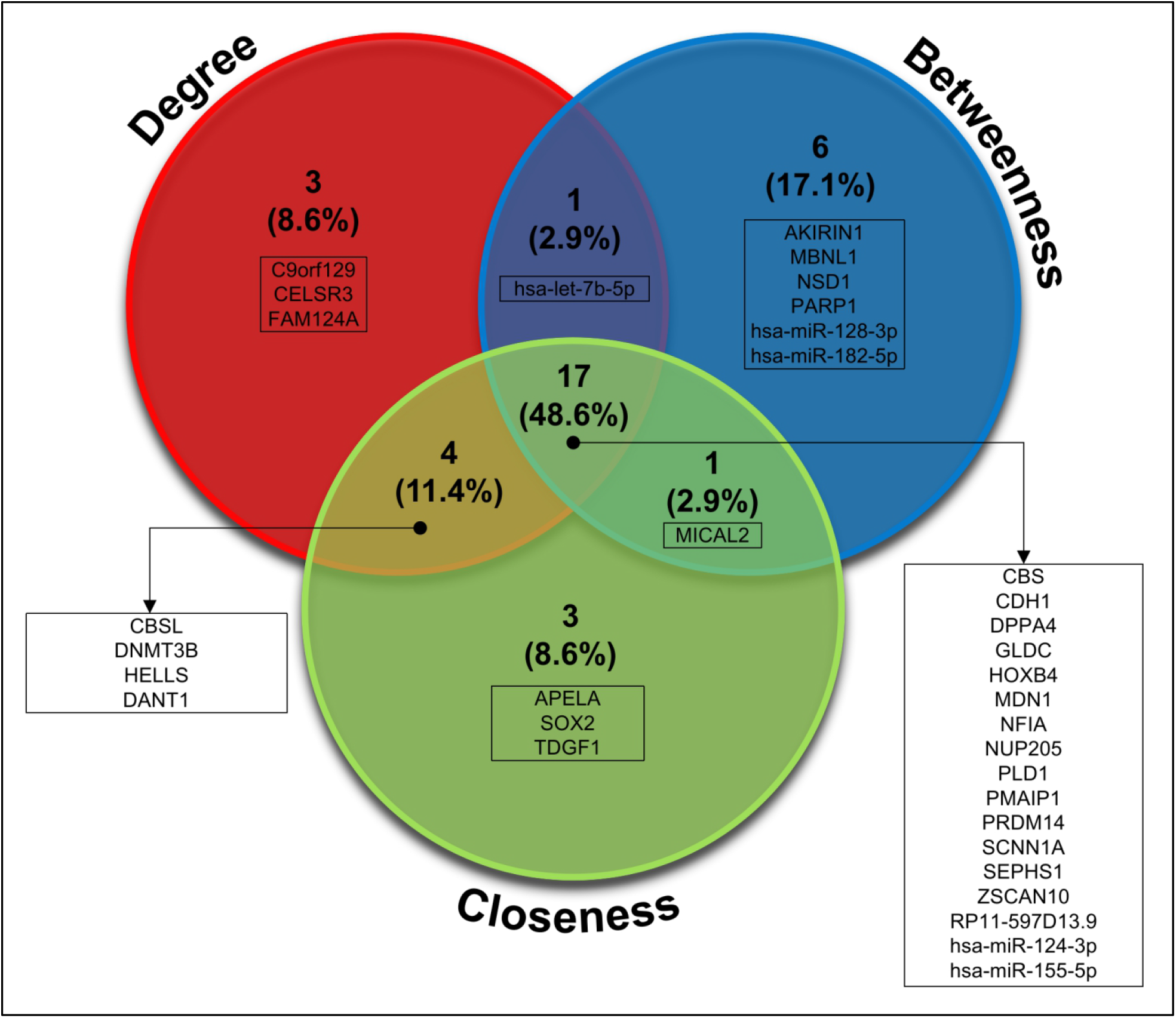
Identification of hub genes by overlapping the top twenty five genes by betweenness centrality, closeness centrality and degree centrality.

**Figure 4:**
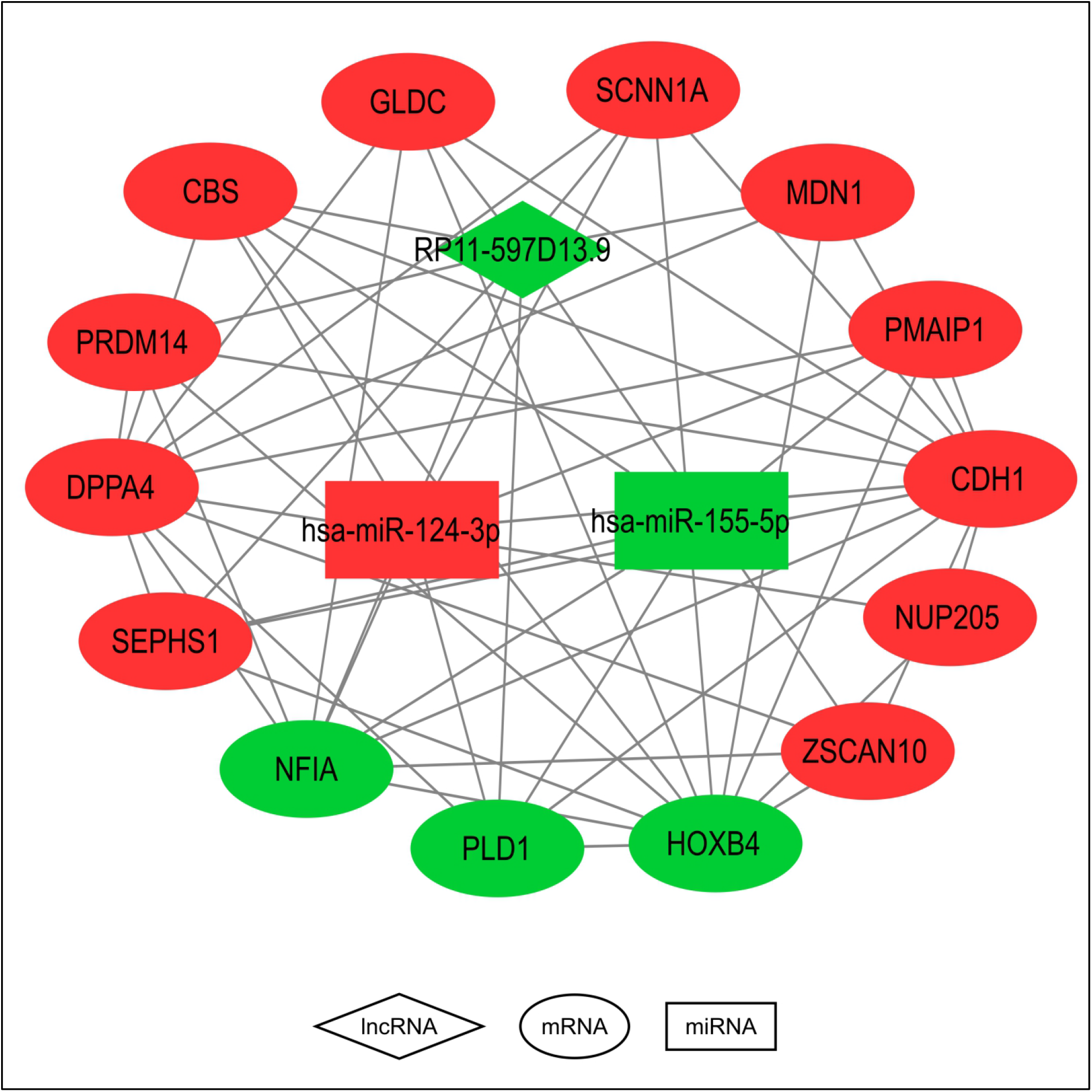
PluriMLMiHubNet – the key MLMi circuitry involved in the establishment and maintenance of pluripotency. The shape of the nodes denotes the gene type – oval for mRNA, diamond for lncRNA and rectangle for miRNA. The upregulated genesin the pluripotent state are coloured red while the downregulated genes are coloured green.

PMAIP1 is essential for cell death in hESCs. It is highly expressed in pluripotent stem cells but decreases during neuronal differentiation and is controlled by POU5F1 (Basundra et al., 2022). SEPHS1 is essential for the survival of hESC. Knockdown of SEPHS1 has been shown to reduce reprogramming efficiency, however, it does not affect the expression of other pluripotency related genes (Lee and Cho, 2019). Unlike the previous few listed genes, NFIA is not expressed in hESCs and its overexpression has been used to facilitate generation of subtype specific astrocytes from human pluripotent stem cells (Li et al., 2018). Similarly, PLD1 is crucial for neurogenesis and is involved in regulation of differentiation of neural stem cells to neurons (Park and Han, 2018). The transcription factor HOXB4 is a master regulator of hematopoiesis and controls several hematopoietic transcription factors and chromatin-modification enzymes. Differentiation of ESCs to hematopoietic cells can be mediated by overexpression of HOXB4 (Fan et al., 2012). The miRNA hsa-miR-124-3p is known to be expressed in undifferentiated hESCs and regulated by one of the key pluripotency related transcription factors POU5F1. Downregulation of this miRNA facilitates migratory cell behaviour (Lee et al., 2010). On the other hand, expression of the miRNA hsa-miR-155-5p has been reported to be associated with suppression of neural stem cell (NSC) self-renewal and promotion of cell differentiation (Teramura and Onodera, 2018). Given the importance of these genes, we believe the seventeen genes must form the key MLMi circuitry in the establishment and maintenance of pluripotency.

### 3.5. Naive and primed state specific MLMi sub-networks

Human pluripotency occurs in two distinct states – naive and primed. While naive pluripotency is the earlier stage of embryonic development, the primed pluripotency occurs during the later stage. In one of our previous work, we had identified the protein coding genes that are key to the naive and primed states of human pluripotency (Supplementary File 4 in Ghosh and Som, 2021). We use this gene list containing 479 naive state specific genes and 507 primed state specific genes to highlight the parts of the PluriMLMiNet that might be critical for the induction and maintenance of the two states. Among the genes overexpressed during pluripotency (i.e., upregulated genes) in the PluriMLMiNet, we found eight genes to be associated with the naive state while 29 genes are associated with the primed state. The first neighbour (nodes directly connected to the selected nodes) lncRNAs and miRNAs of these genes were then retrieved to get the MLMi circuitry of naive and primed state of pluripotency (Figure 5). In the naive state MLMi network (referred as NaiveMLMiNet), the top five mRNAs with the highest degree include RPS6KA1, ZYG11A, ZNF695, ZNF273, and NLRP2, while in the primed state MLMi network (referred as PrimedMLMiNet), the most connected mRNAs are RAB34, TMEM178B, PTPRZ1, USP44, KIF1A, SHISA9, and LRRN1. We observed that each of these genes were not only highly connected in their respective networks, but they also had at least two fold higher expression (log_2_FC > 2) in pluripotent samples as compared to non-pluripotent samples. Thus, these mRNAs have the potential to not only distinguish pluripotent samples from non-pluripotent samples but also to define the state in which the pluripotent samples are in. Apart from the mRNAs, the antisense lncRNA DANT1 was found to be pivotal as it occurred among the most central nodes (by degree, betweenness and closeness centralities) in both the NaiveMLMiNet and PrimedMLMiNet. DANT1 can be seen interacting with crucial naive state markers NLRP2, ZNF695 and ZYG11A, and crucial primed state markers including SALL2, PTPRZ1, OTX2, KIF1A and VRTN (Figure 5 and Supplementary File 3). Among these, we had previously reported ZNF695, SALL2 and OTX2 as crucial transcription factors that might play a switch on-off mechanism in induction of the pluripotent states (Ghosh and Som, 2021). Interestingly, no known interactions were found between DANT1 with any miRNAs, but the mRNAs interacting with DANT1 were found to be regulated by several miRNAs. In the NaiveMLMiNet, the miRNA hsa-miR-30c-2-3p was found to be particularly interesting as it regulated two of the three mRNAs – ZNF695 and ZYG11A – and taken together with DANT1 forms a quartet loop (Figure 6a and Supplementary File 3). On the other hand, in PrimedMLMiNet, we found 17 miRNAs that regulated at least two DANT1 interacting mRNAs (Figure 6b and Supplementary File 3). The miRNA hsa-miR-124-3p controlled at most six of these mRNAs including LRRN1, PTPRZ1, SALL2, SNURF, TEX15, and TOX3. As several of the identified miRNAs were downregulated in pluripotency while the lncRNA DANT1 and the interacting mRNAs were upregulated, it suggests a possible ceRNA mechanism involved in the formation of naive and primed pluripotent states.

**Figure 5:**
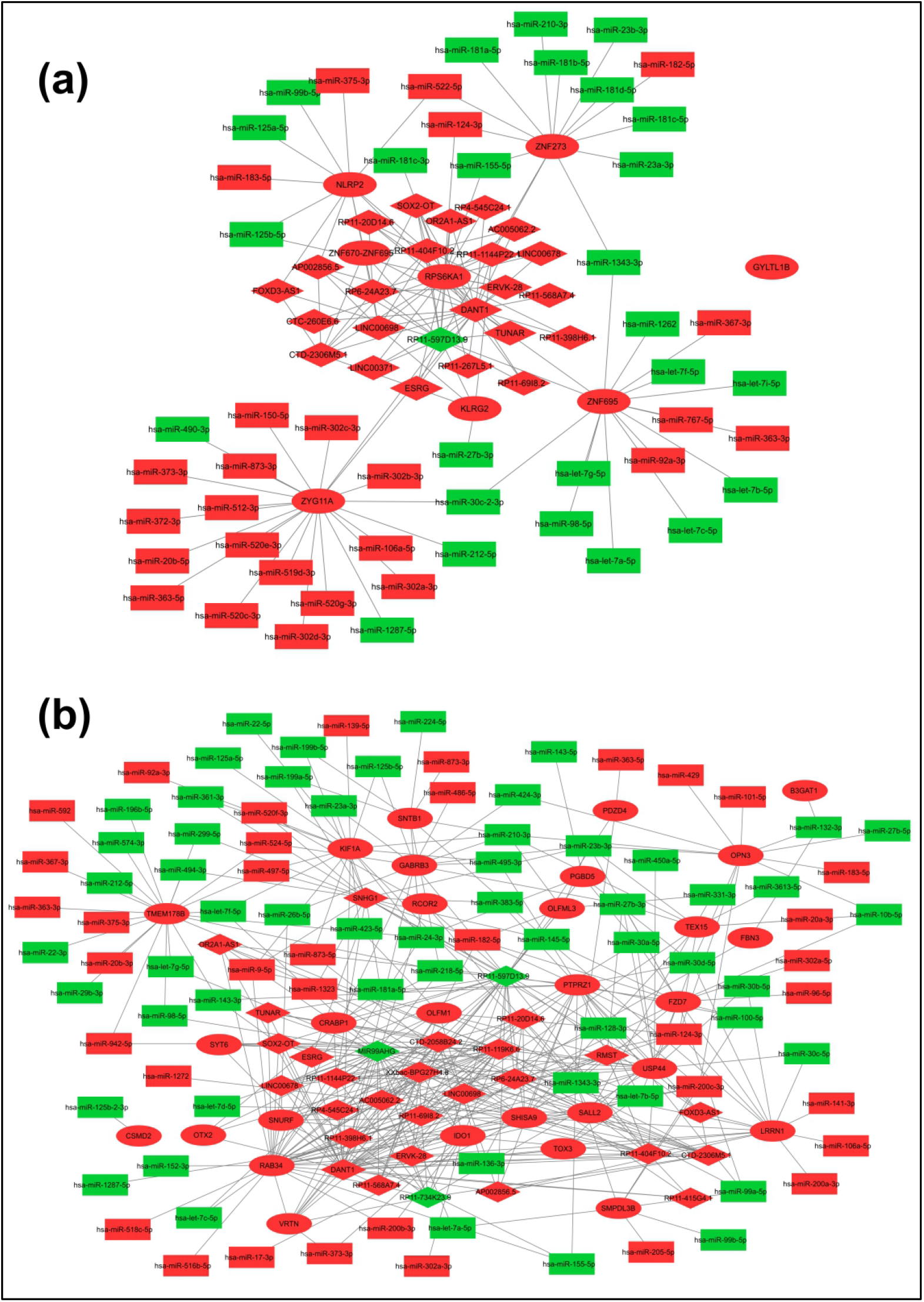
(a) NaiveMLMiNet and (b) PrimedMLMiNet. The networks were extracted by first identifying the naive and primed states specific protein coding genes within the PluriMLMiNet and then retrieving the first neighbour or the immediately connected lncRNAs and miRNAs. The shape of the nodes denotes the gene type – oval for mRNA, diamond for lncRNA and rectangle for miRNA. The upregulated genes in the pluripotent state are coloured red while the downregulated genes are coloured green.

**Figure 6:**
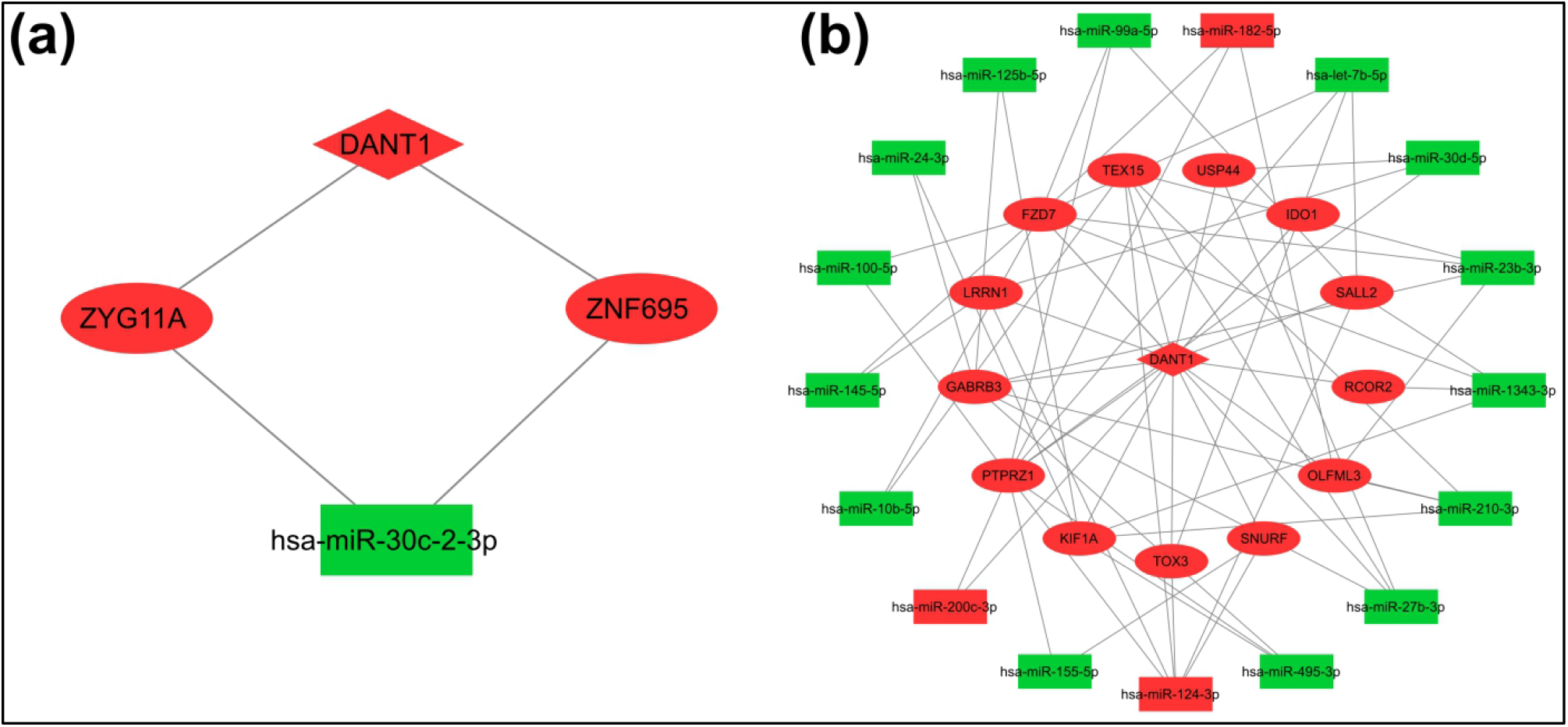
Putative ceRNA network involving DANT1 in (a) naive and (b) primed state. The miRNAs that interact with at least two DANT1 interacting mRNA are shown. The shape of the nodes denotes the gene type – oval for mRNA, diamond for lncRNA and rectangle for miRNA. The upregulated genes in the pluripotent state are coloured red while the downregulated genes are coloured green.

### 3.6. Computational validation of miRNA binding sites in DANT1

To further understand if the lncRNA DANT1 can actually participate to form a ceRNA, we predicted the binding sites within DANT1 for the 18 miRNAs that were found to regulate at least two of the DANT1 interacting mRNAs. Twenty three binding sites and corresponding heteroduplexes were identified (Table 3). However, only three heteroduplexes formed with hsa-miR-30c-2-3p, hsa-miR-210-3p and hsa-let-7b-5p were found to be significant. As mentioned in the previous section, the miRNA hsa-miR-30c-2-3p was found to regulate two DANT1 interacting mRNAs in the NaiveMLMiNet. On the other hand, two distinct sets of DANT1 interacting mRNAs in the PrimedMLMiNet were regulated by the miRNAs hsa-miR-210-3p and hsa-let-7b-5p. The mRNAs GABRB5, KIF1A, OLFML3, and TEX15 were under the control of hsa-miR-210-3p, and FZD7, PTPRZ1, SALL2, and TOX3 were regulated by hsa-let-7b-5p. While DANT1 was found to be upregulated in pluripotent samples as compared to non-pluripotent samples, all the three miRNAs were downregulated. This further confirms our hypothesis that DANT1 along with hsa-miR-30c-2-3p, hsa-miR-210-3p and hsa-let-7b-5p might be forming a ceRNA circuit to regulate several key pluripotency genes.

**Table 3:** List of miRNA-lncRNA heteroduplexes. The significant heteroduplexes (p-value < 0.01) are marked with asterisks (*).

| miR Name | lncRNA | Leftmost binding site position | Folding energy (kcal/mol) | p-value |
| --- | --- | --- | --- | --- |
| hsa-miR-30c-2-3p | DANT1 | 3932 | -18.3 | 0.00139* |
| hsa-miR-210-3p | DANT1 | 1866 | -17 | 0.00644* |
| hsa-let-7b-5p | DANT1 | 1168 | -14.6 | 0.00905* |
| hsa-miR-1343-3p | DANT1 | 696 | -13.6 | 0.0193 |
| hsa-let-7b-5p | DANT1 | 2927 | -17 | 0.0244 |
| hsa-miR-182-5p | DANT1 | 970 | -12.6 | 0.035 |
| hsa-miR-30c-2-3p | DANT1 | 1884 | -16.1 | 0.0503 |
| hsa-miR-1343-3p | DANT1 | 4027 | -15.1 | 0.0618 |
| hsa-let-7b-5p | DANT1 | 1074 | -13.1 | 0.0717 |
| hsa-miR-30c-2-3p | DANT1 | 4192 | -13.6 | 0.0848 |
| hsa-let-7b-5p | DANT1 | 519 | -17.6 | 0.0959 |
| hsa-miR-125b-5p | DANT1 | 609 | -15.7 | 0.1 |
| hsa-miR-1343-3p | DANT1 | 609 | -17.6 | 0.1 |
| hsa-miR-124-3p | DANT1 | 1508 | -14.6 | 0.144 |
| hsa-let-7b-5p | DANT1 | 1478 | -15.7 | 0.144 |
| hsa-miR-27b-3p | DANT1 | 2027 | -14.5 | 0.144 |
| hsa-miR-145-5p | DANT1 | 2034 | -16.3 | 0.144 |
| hsa-miR-30c-2-3p | DANT1 | 3230 | -14.2 | 0.161 |
| hsa-let-7b-5p | DANT1 | 2993 | -13.7 | 0.281 |
| hsa-miR-182-5p | DANT1 | 1437 | -14.3 | 0.315 |
| hsa-miR-210-3p | DANT1 | 1444 | -21.7 | 0.315 |
| hsa-miR-182-5p | DANT1 | 4103 | -18.3 | 0.37 |
| hsa-miR-24-3p | DANT1 | 4324 | -13.3 | 0.384 |

## 4. CONCLUSIONS

The emergence of the role of ncRNAs in biological systems is being increasingly realised. Using a network biology approach on transcriptomic data, we attempted to uncover the mRNA-lncRNA-miRNA interaction landscape in human pluripotency. We reconstructed the core MLMi network – PluriMLMiNet – that we believe must be crucial for the induction and maintenance of pluripotency in humans. Interestingly, we observed that the hub of PluriMLMiNet (PluriMLMiHubNet) did not include some of the most popular pluripotency genes – NANOG, POU5F1 and SOX2 – suggesting a possible alternate circuit for maintenance of pluripotency. Further, based on the pluripotency state specific gene lists from our previous finding, we also extracted the naive and primed state specific MLMi networks, NaiveMLMiNet and PrimedMLMiNet respectively. Our findings open up a new set of marker genes (RPS6KA1, ZYG11A, ZNF695, ZNF273, and NLRP2 for naive state, and RAB34, TMEM178B, PTPRZ1, USP44, KIF1A and LRRN1 for primed state) that can not only be used to distinguish pluripotent state from non-pluripotent state but also identify the intra-pluripotency state it exists in. Mechanistically, we identified the lncRNA DANT1 to be crucial as it was found to be forming a bridge between the naive and primed state specific MLMi networks. Also, the pattern of its interactions with the mRNAs and miRNAs suggests its possible involvement in the ceRNA mechanism. We computationally confirmed that DANT1 harbours binding sites for the miRNAs hsa-miR-30c-2-3p, hsa-miR-210-3p and hsa-let-7b-5p. Thus, our work opens up newer perspectives into the mechanism of pluripotency in humans.

## ABBREVIATIONS

ceRNA: Competing Endogenous Ribonucleic Acid
ENA: European Nucleotide Archive
ESC: Embryonic Stem Cell
lncRNA: long non-coding Ribonucleic Acid
MLMi: mRNA-lncRNA-miRNA
mRNA: Messenger Ribonucleic Acid
miRNA: Micro Ribonucleic Acid
TOM: Topological Overlap Measure
WGCNA: Weighted Gene Co-expression Network Analysis

## ACKNOWLEDGEMENTS

AS thank the Department of Biotechnology (DBT) for providing financial assistance for the project.

## LEGENDS

**Supplementary File 1:** Details of raw RNA-seq and miRNA-seq data included in the study.

**Supplementary File 2:** Results of differential gene expression analysis.

**Supplementary File 3:** Interactive cytoscape file containing the PluriMLMiNet, PluriMLMiHubNet, NaiveMLMiNet and PrimedMLMiNet.

